# Brain Charts for People Living with Multiple Sclerosis

**DOI:** 10.1101/2023.11.02.565251

**Authors:** A. Keshavan, D. Peterson, A. Alexander-Bloch, K.M. Leyden

## Abstract

Multiple sclerosis (MS) is associated with brain volume loss throughout the disease course. Currently available automated segmentation methods can measure total brain volume as well as ventricular volume, which has been advocated as a robust surrogate for brain volume based on clinically acquired magnetic resonance imaging (MRI). However, brain and ventricle volumes change naturally with age and may be susceptible to biases from differences in acquisition hardware, imaging protocol, and image quality, in addition to statistical biases such as regression to the mean. In this work, brain charts for people living with MS were established that account for patient biological sex, age, and differences in acquisition and image quality. 379 subjects were imaged longitudinally at 5 MS centers using 13 MRI scanner models from 2 scanner manufacturers employing a variety of protocols that included T1-weighted and T2-weighted FLAIR imaging. Generalized Additive Models for Location Scale and Shape (GAMLSS) were employed, and scanner metadata as well as automated assessments of image quality were modeled. Cross-sectional brain charts and conditional, longitudinal brain charts were estimated separately in female and male participants resulting in interpretable and intuitive centile estimates. These findings indicate that brain charts for people living with MS are a promising method for turning quantitative volumetrics into actionable knowledge about a patient’s disease.

**Highlights:**

A. Interpreting observed changes in brain volume can be challenging due to statistical biases including regression to the mean.
B. Brain size changes naturally with age and may be susceptible to biases associated with acquisition hardware, imaging protocol, and image quality.
C. Brain charts for people living with MS are a promising method for translating quantitative volumetrics into interpretable knowledge about a patient’s disease.

## Introduction

Improving brain MRI-based prediction of clinical outcomes for people living with neurological conditions such as multiple sclerosis (MS) may depend on the ability to accurately and sensitively quantify an individual’s pathology relative to typical disease progression. Progression of MS pathology includes not only pathognomic white matter lesions but also neurodegeneration that manifests as decreased brain volume and increased ventricular volume (Filippi et al., 2016; Klaver et al., 2013; Lassmann, 2018). MRI measurements of tissue volume and longitudinal changes in MRI volumetric measurements over time have been associated with cognitive impairment (Eshaghi et al., 2018; Preziosa et al., 2016) and physical disability (Matthews et al., 2023). Individuals with specific patterns of deviation relative to typical disease progression may have more or less severe forms of disease requiring differential clinical management. However, the interpretation of quantitative assessments of brain volume in people living with MS can be challenging due to confounding with sex differences, age-expected changes, and other demographic factors. In this context, normative models of brain MRI data may increase sensitivity to identify clinically-relevant alterations in individual patients (Marquand et al., 2019).

Normative modeling is an increasingly popular statistical framework to quantify imaging pathology relative to a reference population in human brain MRI research (Bethlehem et al., 2020; Marquand et al., 2016; Wolfers et al., 2018, 2020; Ziegler et al., 2014). Conceptually, the overarching goal of these studies is similar to the use-case of pediatric growth charts for height or weight, which are a critical aspect of clinical care and research, based on statistical models that have evolved since growth charts were first introduced in the 19th century (Cole, 2012; Cole & Green, 1992; Wei et al., 2006). For the development of growth charts, the World Health Organization has previously recommended a distributional regression framework known as generalized additive models for location, scale and shape (GAMLSS) due to its combination of modeling flexibility and statistical rigor (Borghi et al., 2006). GAMLSS allow for the modeling of data whose distribution does not follow an exponential family as in standard generalized additive modeling (Stasinopoulos et al., 2018). Furthermore, this approach allows for modeling the mean structure as well as the variance, skewness, and kurtosis in terms of flexible nonlinear associations with covariates of interest (Rigby et al., 2019). This modeling approach was recently employed in a large international effort to develop brain charts based on cross-sectional MRI data, which were shown to be highly sensitive to alterations in patients with neurodegenerative disorders and other neuropsychiatric conditions (Bethlehem et al., 2022).

Neuroimaging applications have largely focused on brain charts based on cross-sectional reference samples, which can be applied post hoc to longitudinal data. However, there are clear advantages to explicitly deriving models based on a longitudinal reference sample. Within-individual change can then be quantified using longitudinal growth charts, which model interindividual variability in velocity rather than distance, and can also account for the expected regression to the mean of extreme measurements over time (Cole, 1994; Wei & He, 2006). Within-individual change can be formalized in GAMLSS models by conditioning data from one time point on data obtained at a previous time point. Such models are particularly compelling for research or clinical contexts where longitudinal data is the norm rather than the exception, as is the case for individuals living with MS. As is true for pediatric growth charts (Canadian Paediatric Society, 2010), the degree of within-individual change between measurements may be of clinical concern even if each measurement lies within the “normal” range if considered independently.

For both cross-sectional and longitudinal models, the interpretation of normative models depends on the nature of the reference sample. While most prior work in neuroimaging normative modeling has used “healthy controls” as the reference sample, comparing individuals living with MS to a reference composed of other individuals living with MS may be more informative. In many if not most cases, the diagnosis of MS is not in question, but individuals undergo regular MRI scans to monitor disease progression (Kaunzner & Gauthier, 2017). In this context, while MS is generally expected to be associated with a more pronounced decline in gray matter volume compared to typical aging, individuals with different patterns of disease progression relative to an MS-specific model may represent informative subgroups who benefit from alternative management strategies or experimental therapeutics. The utility of population-specific normative models has been anticipated in other areas of medicine, for instance, the use of specific growth charts for children with known neurogenetic conditions (Cronk et al., 1988; Lyon et al., 1985). Sex-specific MS models are also critical, because not only do average brain volumes differ across the sexes, but men living with MS tend to have accelerated brain volume loss compared to women (Voskuhl et al., 2020).

In addition to measurements of brain parenchymal volume, changes in ventricular volume are sensitive measures of neuropathology. Shrinking of cortical and subcortical gray and white matter, coupled with expansion of the ventricular compartment, is a typical part of human aging (Barron et al., 1976; Bethlehem et al., 2022). This trend is more pronounced in neurodegenerative disease, for example increased volume in the inferior portion of lateral ventricles is a robust marker of tissue loss in medial temporal lobe structures in Alzhiemer’s disease (Brewer et al., 2009; Jack et al., 1997; Murphy et al., 2010). Certain patterns of ventricular volume changes have also been specifically linked to MS (Brex et al., 2000; Dalton et al., 2006; Millward et al., 2020; Sinnecker et al., 2020), resulting in an outstanding clinical and scientific need for brain charts of MRI measurements of both total brain volume and ventricular volume in MS.

The present study extends prior research in MS and normative modeling in several novel directions. Using GAMLSS, normative models of total brain volume and ventricular volume in MS (“MS brain charts”) were derived using a large multi-site dataset that includes 13 MRI scanners and 2 MRI scanner manufacturers. To maximize clinical applicability, the FDA-cleared NeuroQuant software tool was applied for tissue segmentation (NeuroQuant MS, v3.1, Cortechs.ai), and an automated measure of image quality was incorporated into brain chart models. Distinct brain chart models were estimated for cross-sectional evaluation and longitudinal monitoring. We hypothesized that MS brain charts would provide a personalized measure with improved sensitivity to interindividual differences in brain atrophy for people living with MS.

## Methods

### Experimental Methods

Real-world MRI data were aggregated from 5 MS centers using 13 MRI scanner models from 2 scanner manufacturers employing a variety of standard-of-care protocols that included T1-weighted and T2-weighted FLAIR imaging.

Total brain volume and total ventricle volume were extracted using the FDA-cleared NeuroQuant software tool (NeuroQuant MS, v3.1, Cortechs.ai), and automated image quality assessment was employed using the MRIqc tool (Esteban et al., 2017).

To assess the distribution of total brain volume and ventricular volume through the age span and build MS brain charts, GAMLSS [2] were employed for each measurement individually. GAMLSS allow for the modeling of data whose distribution does not follow an exponential family as in standard generalized additive modeling. The GAMLSS approach employs the specification of a 3- or 4-parameter model to link the mean, variance, skewness, and kurtosis to the predictor variables (Bethlehem et al., 2022). As GAMLSS are complex models, convergence in limited sample sizes such as that under study here is not guaranteed. To simplify model-building, cross-sectional modeling was conducted in the last measurement from each subject for cross-sectional models and the last pair of measurements for longitudinal models. Subsequent to model building, predictors of volume distribution were assessed using Wald tests (Wald, 1943) with no corrections for multiple comparisons due to the exploratory nature of the study.

### Cross-Sectional Brain Chart Development

Denote the volume measurement under study for subject 𝑖 at visit 𝑗 by 𝑌_𝑖𝑗_, their age by 𝐴_𝑖𝑗_, and additional covariates of interest (see below) by 𝑋_𝑖𝑗_. For each parameter θ_𝑘_ (θ_𝑘_ = µ for 𝑘 = 1, for example) to be modeled, further define an appropriate link function 𝑔_𝑘_ and a family for the distribution of 𝑌_𝑖𝑗_. The GAMLSS model employed was

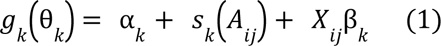

for 𝑘 = 1, …, 3 or 4, depending on the choice of distributional family. Note that this is a simplification of the general GAMLSS model without a random effect structure in the mean specification, and that 𝑠_𝑘_(𝐴_𝑖𝑗_) denotes an estimated smooth function of 𝐴_𝑖𝑗_ for each 𝑘.

Separate cross-sectional models were employed for biological males and females. Employing worm plots (van Buuren, 2007) and visual assessment of fitted centiles, model building and checking was conducted for each sex. Variables considered in the modeling process were age, contrast-to-noise ratio (CNR, from MRIqc applied to the T1-weighted imaging), scanner manufacturer, and scanner field strength. Scanner serial number, model, and acquisition site were not included in modeling as the goal was to maximize generalizability. Age was modeled using a penalized beta spline, which allows for flexible nonlinear associations while mitigating roughness and overfitting. The default three-parameter Box-Cox-Cole-Green distribution was employed, and more complex distributional families were considered if worm plot analysis indicated insufficient model flexibility. Model building was conducted by sequential inspection of worm plots starting with mean modeling and then subsequently for higher-order models considering variance, skewness, and finally kurtosis. Predictors for which worm plot analysis indicated an association for a higher-order moment were appropriately included in lower-order moment models. If model fit for a higher-order model was observed to be inferior to a lower-order model, the lower-order model was selected. See Supplementary Figure 1 for an illustrative example of a worm plot.

### Longitudinal Brain Chart Development

While a patient’s current brain volume centile is informative, the availability of longitudinal data allows for more granular assessments. Indeed, a participant whose brain volume is near the population median might change substantially relative to previous measurements but still have near-median brain volume compared with other participants. Brain charts based only on cross-sectional data have also been reported to systematically underestimate brain changes directly measured in longitudinal data (Biase et al., 2023). This emphasizes the importance of longitudinal brain charts, in which modeling is conducted conditionally on previous measurements (see Figure 3, inspired by (Cole, 1994)).

To model longitudinal measurements, the GAMLSS approach described in equation (1) was employed with a parametric form determined by the model-building process. In particular, and motivated by (Cole, 1994; Wei et al., 2006; Wei & He, 2006):

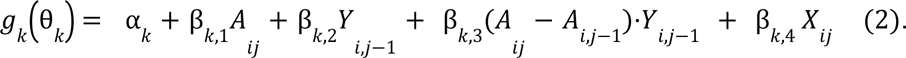

Covariates 𝑋𝑖𝑗 were considered at the time of the follow-up visit 𝑗 for the longitudinal setting. Each term was examined for improvement in fit based on worm plot analysis, and the distributional family was selected similarly to the cross-sectional modeling case.

## Results

### Sample Description

758 brain MRIs were acquired in 376 people living with MS (see Table 1). 262 images were acquired at 1.5T and 496 images employed 3T MRI equipment. 49% of images were acquired on scanners manufactured by General Electric (GE, Milwaukee, WI) and 51% were acquired on Siemens scanners (Erlangen, Germany). The sample was 76.5% female, and the median age was 43.2 years (range 20 – 76). Cross-sectional modeling was conducted in 379 MRIs, and longitudinal modeling was conducted in 558 MRI scans (279 pairs, see Figure 1). All images were employed in subsequent model diagnostics and visualizations.

**Figure 1:**
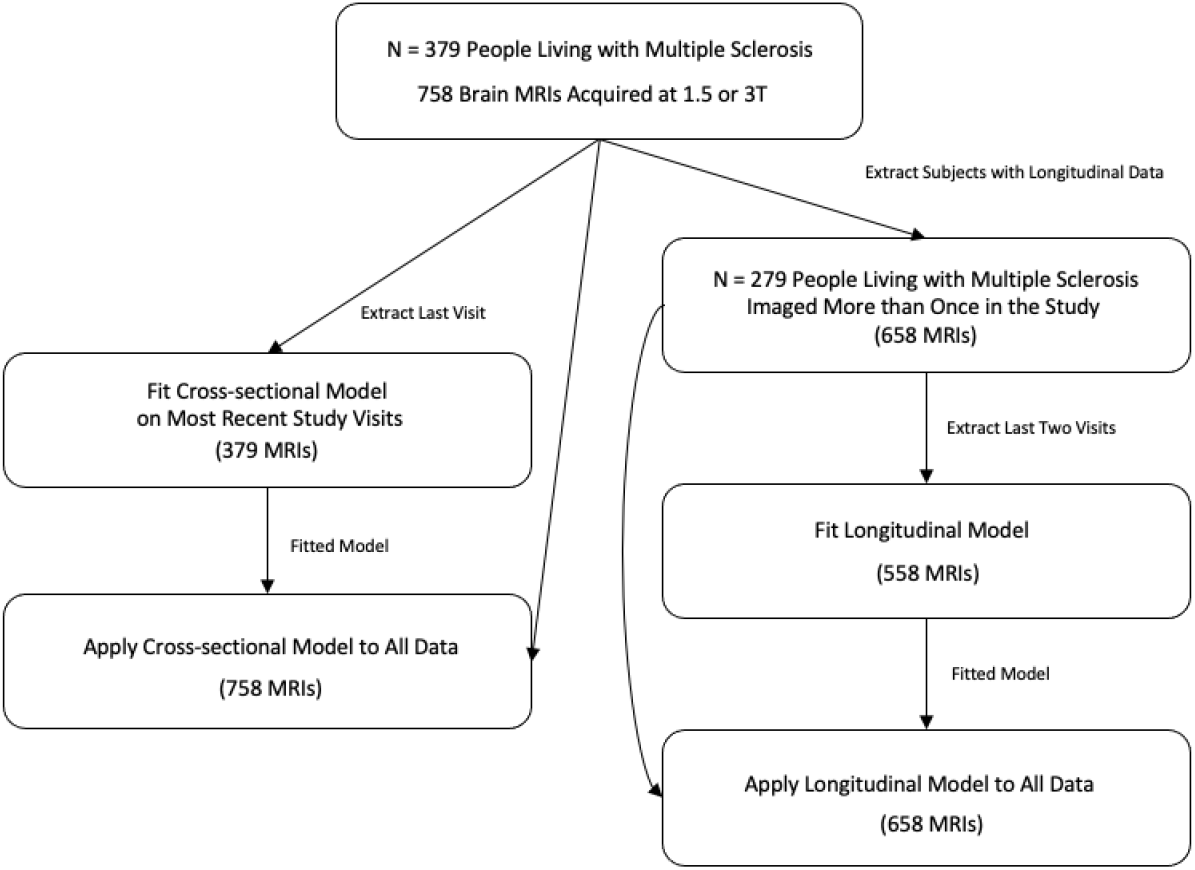
Flowchart demonstrating the dataset under study and modeling strategy.

**Table 1:** Demographics and summary information for the study sample.

|  | Female<br>(N=290) | Male<br>(N=89) | Overall<br>(N=379) |
| --- | --- | --- | --- |
| **Scanner Manufacturer (Baseline)** |  |  |  |
| GE MEDICAL SYSTEMS | 137 (47.2%) | 48 (53.9%) | 185 (48.8%) |
| SIEMENS | 153 (52.8%) | 41 (46.1%) | 194 (51.2%) |
| **Age (years)** |  |  |  |
| Mean (SD) | 43.3 (12.7) | 43.4 (11.9) | 43.3 (12.5) |
| Median [Min, Max] | 42.0 [20.0, 76.0] | 44.0 [20.0, 67.0] | 42.0 [20.0, 76.0] |
| **T1-weighted Contrast to Noise Ratio** |  |  |  |
| Mean (SD) | 2.99 (0.598) | 2.77 (0.539) | 2.94 (0.591) |
| Median [Min, Max] | 3.03 [0.553, 4.41] | 2.72 [1.73, 4.30] | 2.99 [0.553, 4.41] |
| **Baseline Total Brain Volume (mL)** |  |  |  |
| Mean (SD) | 1120 (119) | 1270 (118) | 1160 (134) |
| Median [Min, Max] | 1120 [800, 1420] | 1270 [977, 1550] | 1160 [800, 1550] |
| **Baseline Ventricle Volume (mL)** |  |  |  |
| Mean (SD) | 31.7 (17.6) | 39.3 (17.2) | 33.5 (17.8) |
| Median [Min, Max] | 26.6 [10.2, 133] | 34.5 [14.1, 101] | 28.9 [10.2, 133] |
| **Acquisition** |  |  |  |
| Site 1 | 66 (22.8%) | 20 (22.5%) | 86 (22.7%) |
| Site 2 | 67 (23.1%) | 20 (22.5%) | 87 (23.0%) |
| Site 3 | 6 (2.1%) | 1 (1.1%) | 7 (1.8%) |
| Site 4 | 78 (26.9%) | 23 (25.8%) | 101 (26.6%) |
| Site 5 | 73 (25.2%) | 25 (28.1%) | 98 (25.9%) |

### Model Fitting

Cross-sectional GAMLSS modeling of total brain volume and ventricle volume for males and females was conducted separately based on model (1) (see Figure 2). Worm plots indicated nonlinear age associations as well as linear contributions of T1 CNR, field strength, and scanner manufacturer for the mean and variance models for both sexes (see Supplementary Figure 1). In the larger female sample, skewness was also noted for total brain volume and a nonlinear age term and linear terms for T1 CNR and scanner manufacturer were included. For ventricle volume in females, skewness was observed and these same terms as well as field strength were included.

**Figure 2:**
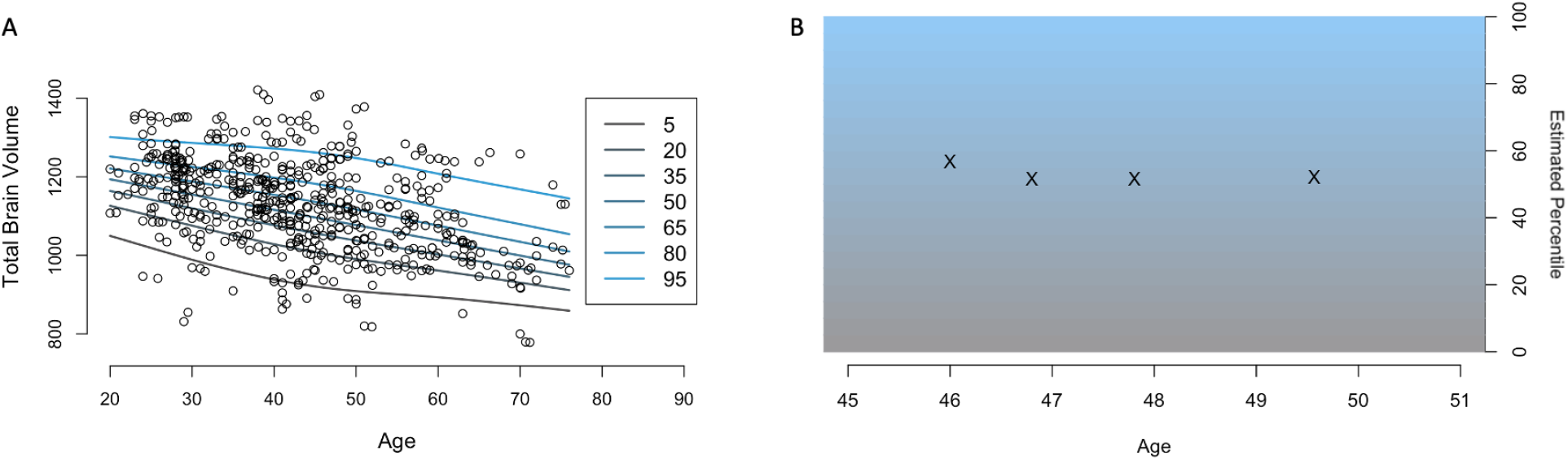
Demonstration of growth chart methodology. Panel A (left) shows n=573 total brain volume measurements in mL from the female participants in our study (n=290) in circles through the age span. Lines indicate the fitted centiles for expected volume at each age, with different colors indicating example percentiles. Panel B (right) shows one particular subject’s cross-sectional percentiles as she ages. At age 46, her brain is estimated to be around the 57th percentile; however, at ages 47, 48, and 50 her brain appears to be very close to the median, at the 52nd percentile.

**Figure 3:**
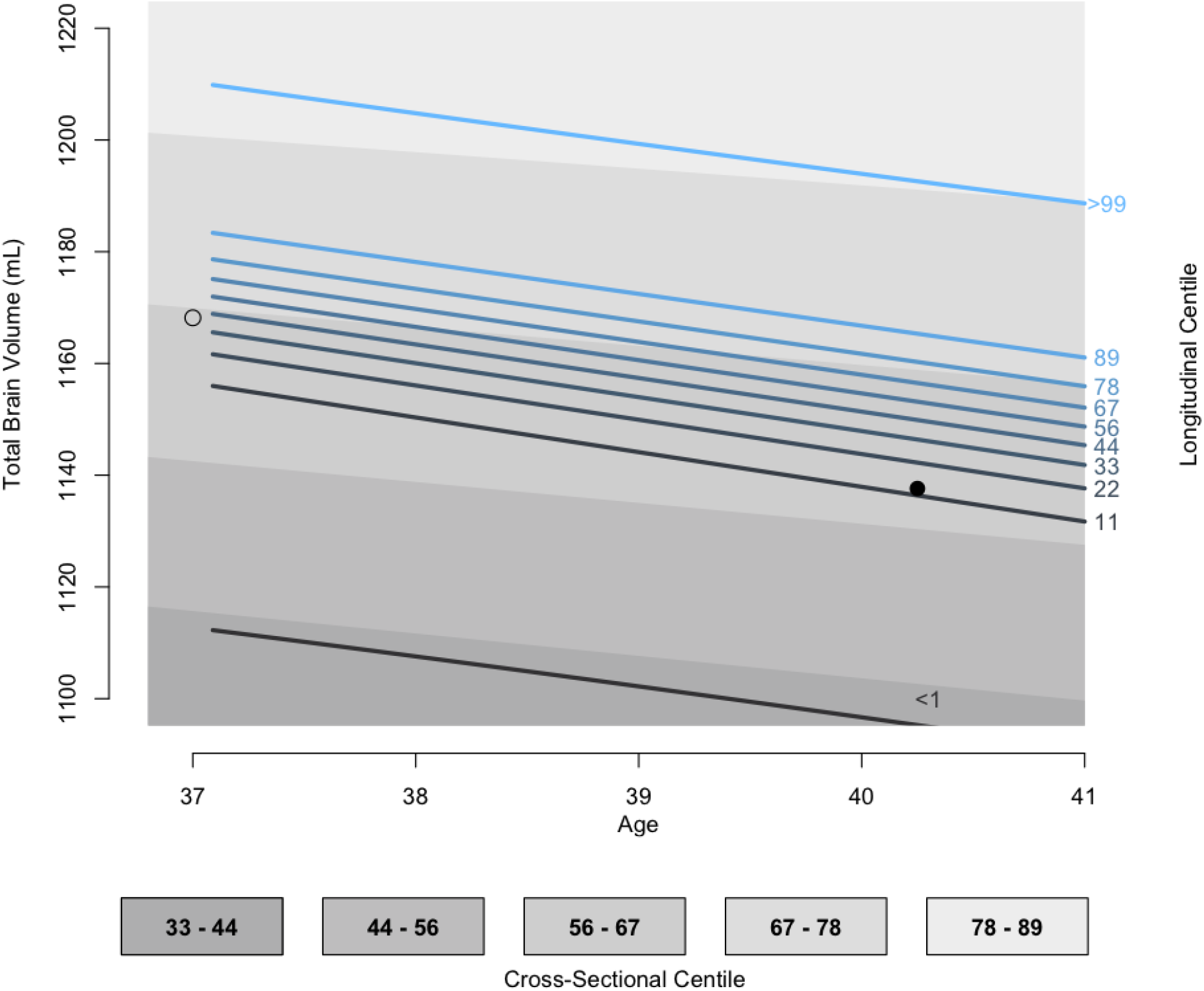
Example demonstrating the important distinction between cross-sectional and longitudinal growth charts. Background colors show cross-sectional centiles, and lines indicating longitudinal centiles as labeled to the right. Total brain volume at age 37 for an example subject is shown in the empty circle at 1168mL, which corresponds to above the median in this study. At age 40, her brain volume decreased by 31 mL which was still above the median when viewed as a single cross-sectional measurement. However, when viewed in light of the measurement at age 37, this is in fact at the 14th percentile of expected brain volume, indicating a potentially clinically concerning decrease. Inspired by Cole (1994).

Worm plot analysis of longitudinal GAMLSS modeling of total brain volume based on model (2) indicated linear mean associations with T1 CNR (at follow-up), scanner manufacturer, and field strength in addition to the first four terms in model (2). Variance associations with those same predictors were also observed. Worm plots indicated models for skewness and kurtosis in females and the Box-Cox *t (Rigby & Stasinopoulos, 2006)* distribution was employed with a nonlinear age term, scanner manufacturer, field strength, and T1 CNR for both ventricle volume as well as total brain volume.

### Predictors of Volume Distribution

GAMLSS-based fixed effect inference was conducted on all non-age-related terms to assess which factors were significantly associated with the cross-sectional and longitudinal distribution of the modeled volumes.

Cross-sectionally, Siemens scanners showed lower average brain volume in males (57mL, p<0.05). No statistically significant associations were observed between acquisition parameters and total brain volume in females. Longitudinally, average brain volume was slightly lower on Siemens scans (10mL, p<0.04), and greater brain volume variance was associated with GE scanners (p<0.03) in males. In females, 3T magnets were associated with smaller variance (p<0.02) and Siemens scanners (p<0.001).

For ventricle volume, cross-sectional GAMLSS modeling in males found higher average volume on Siemens scanners (6mL, p<0.04) and lower variance with higher T1 CNR (p<0.02) and GE scanners (p<0.04). No statistically significant associations were observed between acquisition parameters and ventricle volume in females. Longitudinal modeling in males indicated increased ventricle volume was associated with lower T1 CNR (p<0.001) and Siemens scanners (0.4mL, p<0.001). 1.5T imaging, GE scanners, and higher T1 CNR were associated with smaller variance in ventricle volume in males (all p<0.001). On the other hand, larger follow-up ventricle volume was associated with 1.5T imaging (p<0.05) in females. Smaller variance was associated with higher T1 CNR (p<0.02) and 3T imaging (p<0.01). T1 CNR was associated with greater skewness and less kurtosis (both p<0.001), and Siemens scanners exhibited more kurtosis (p<0.02).

### Visualizing and Interpreting Brain Chart

Figure 4 shows examples of fitted cross-sectional and longitudinal models in three subjects labeled as A, B, and C.

**Figure 4:**
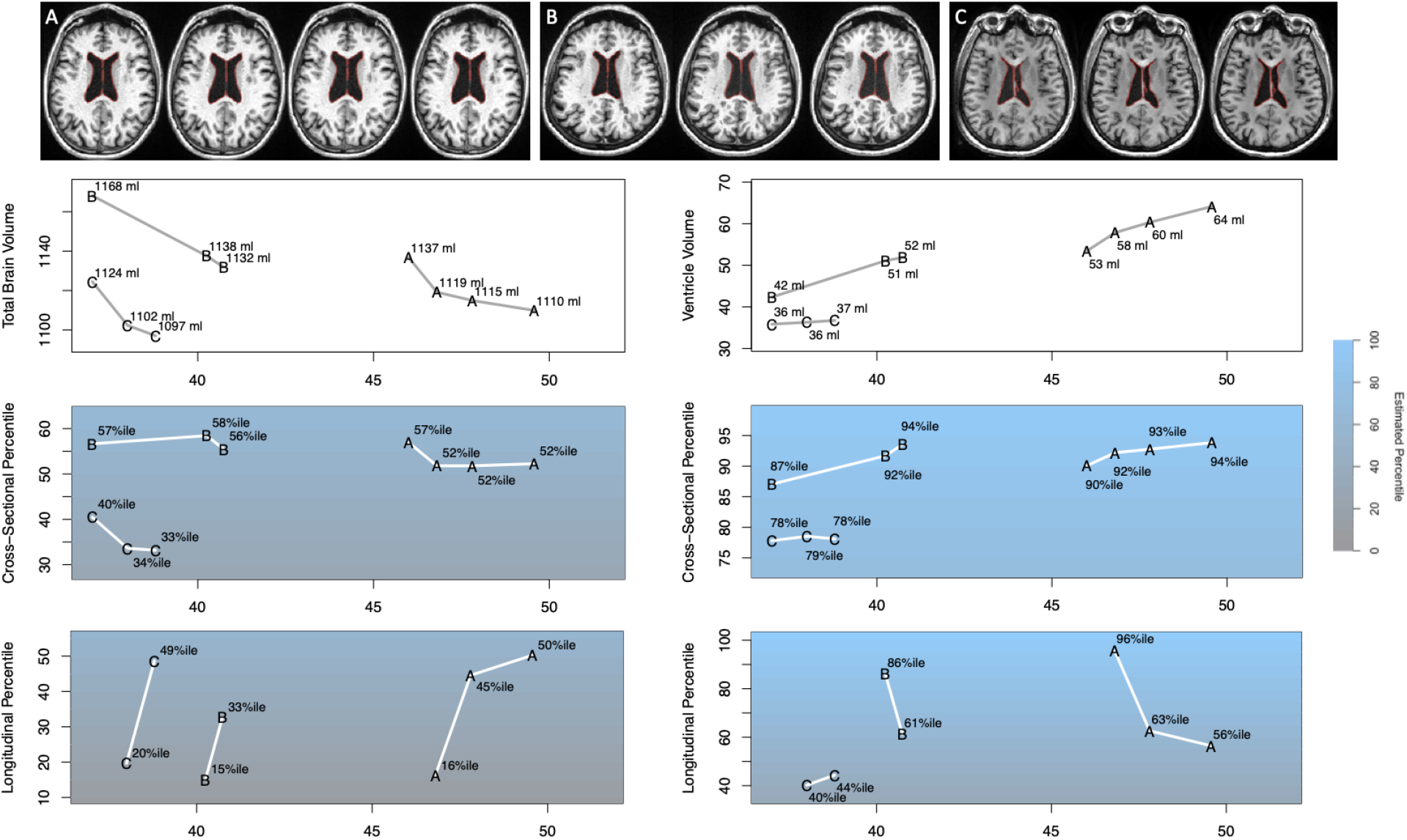
Results from cross-sectional and longitudinal GAMLSS modeling in three subjects labeled A, B, and C. The first row shows axial views of each T1-weighted image with a red outline around the ventricles. Visual inspection shows subtle brain volume loss in all three cases, as expected in MS. The lower section of the figure shows, in rows, raw brain volumes, cross-sectional percentiles, and longitudinal percentiles with total brain volume in the left column and ventricle volume in the right column. Note that the bottom row shows one fewer point for each subject compared with the second-to-last row, as no previous data were available to assess longitudinal change for the baseline scans. In all three subjects, brain volume decreases and ventricle volume increases, but cross-sectional percentiles are somewhat stable. Longitudinal centiles for brain volume are low for each of the second timepoints, but closer to the median subsequently. For ventricle volume, a similar (but opposite direction) trend was observed for subjects A and B, but subject C’s cross-sectional and longitudinal centiles are more stable. This may indicate that the brain volume change observed in subject C was anatomically focused in a brain region or tissue compartment where changes in volume were less associated with corresponding ventricular change.

Subject A was observed at four visits between the ages of 46 and 50, whereas Subject B was observed at three visits between 37 and 41, and Subject C was observed at three visits between 37 and 39. The top row of Figure 4 shows axial views of the T1-weighted imaging; Subject A was imaged on a 1.5T Siemens Avanto scanner at all time points, and Subject C was imaged on two 3T GE scanners. Subject B, on the other hand, was imaged at the first time point on a 1.5T Avanto, and at the following time points on 3T Siemens Skyra Fit equipment.

All three subjects showed decreasing brain volume and increasing ventricle volume during the study follow-up. They also showed a more prominent brain volume drop between the first two timepoints, but none showed a change in cross-sectional MS brain chart percentile of more than 10% (see third row of Figure 4). However, longitudinal centiles for the first follow-up visits for all three subjects were in the bottom quintile indicating a large within-subject relative change. On subsequent follow-up visits, brain volumes and observed decreases therein were within the middle tertile.

A similar but slightly different pattern was observed in ventricle volumes. All three subjects had relatively large ventricles, with Subjects A and B near or in the top decile and Subject C in the top quartile. Subjects A and B also showed a rapid increase in ventricle volume between the first two visits contemporaneous with the brain volume loss described above, with longitudinal centiles also over 85%. However Subject C’s ventricle volume was quite stable, with longitudinal centiles below the median for ventricular growth.

## Discussion

By providing quantitative models of interindividual variability in brain volume measurements and brain volume changes over time, brain charts have the potential to significantly advance our understanding of brain volume alterations associated with MS. The present study uses a real-world multi-center cohort of clinically acquired brain MRI images to develop disease-specific normative models of total brain volume and ventricle volume. A WHO-recommended statistical modeling framework with GAMLSS (Borghi et al., 2006; Stasinopoulos et al., 2018) yielded MS brain charts for males and females and for both cross-sectional and conditional longitudinal settings. Measurements of brain volume were subtly influenced by MRI scanner hardware and MRI scan quality, supporting their inclusion as covariates in brain chart models. Compared to cross-sectional models, longitudinal models yielded more precise and sensitive descriptions of changes in both total brain and ventricle volume. Collectively, these results point to the capacity for MS brain charts to advance imaging biomarker development and to augment the clinical interpretation of brain MRIs for people living with the disease.

It is predicted by prior literature that MS brain charts would benefit from the inclusion of information about MRI scan quality, scanner manufacturer, and acquisition parameters. It is well known that MRI scanner platform can bias automated volumetric measurements derived from brain MRI images (Agartz et al., 2001; Jovicich et al., 2013; Schnack et al., 2004; Shinohara et al., 2017). Systematic differences in estimated brain volumes between scanner manufacturers, as was observed in this study, have been documented extensively (Auzias et al., 2016; Jovicich et al., 2009; Lee et al., 2019). Moreover, statistical harmonization has been demonstrated to mitigate this bias in a variety of contexts (Chen et al., 2022; Fortin et al., 2018). Relatedly, it is known that automated quantitative measures derived from structural brain MRI are sensitive to interindividual differences in MRI scan quality that can result, for instance, from interindividual differences in in-scanner motion (Alexander-Bloch et al., 2016; Blumenthal et al., 2002). It is increasingly common practice in neuroimaging studies for statistical models to include as covariates manual image quality ratings or automated measures related to image quality, such as the Euler number from Freesurfer’s cortical reconstruction or statistics output by the MRIqc tool (Esteban et al., 2017; Rosen et al., 2018). Consistent with this prior literature, conditioning GAMLSS models on scanner platform, field strength, and MRI image quality improved model fit for centiles scores for volumetric measures provided by the present study.

MS brain charts benefited from the inclusion of longitudinal data in statistical models. While cross-sectional models can be utilized to track changes over time, longitudinal models are likely superior for the detection and quantification of “surprising” changes in a measured variable (Cole, 1994; Wei & He, 2006). These changes are of high clinical relevance for people living with MS, as brain volume loss may be associated with disease progression and worsening cognitive disability (De Stefano et al., 2014; Eshaghi et al., 2018; Frau et al., 2018; Sastre-Garriga et al., 2017). In the present study, conditioning GAMLSS models on previous brain volume measurements in the same participant improved the quality of statistical models. When directly comparing cross-sectional and longitudinal MS brain charts, longitudinal models provided increased sensitivity to disease-related change in both total brain volume and ventricle volume.

In presenting MS growth charts for both total brain volume and ventricle volume, we highlight the complementary information provided by different imaging-derived neuroanatomical phenotypes. The relationship between total brain and ventricular volume is nonlinear and age-varying (Bethlehem et al., 2022). In typical aging, total brain volume decreases while ventricular volume increases, a pattern that is accentuated in neurodegenerative disorders including MS (Brex et al., 2000; Dalton et al., 2006; Millward et al., 2020; Sinnecker et al., 2020). Our results demonstrate that measurements of total brain volume versus ventricle volume may have differential sensitivity to disease progression in specific individuals. The differential sensitivity may result from brain volume loss targeting different anatomical foci (Haider et al., 2016), for instance subcortical nuclei or white matter compartments that are expected to have different relationships with ventricular expansion.

There are several limitations of the presented analyses, many of which stem from the limited sample size available. First, both the cross-sectional and longitudinal models were built based on subsets of the data due to issues with convergence when modeling the full longitudinal data. Second, the value of each technical variable for the purpose of improving model fit and accuracy of centile estimation was conducted by testing after model construction. Testing after model construction may be biased due to post-selection inference, although since visual assessments were conducted based on residuals as opposed to stepwise regression using p-values the degree of this bias is expected to be lessened. Furthermore, additional quality predictors would likely be helpful in larger sample size modeling. For example, baseline quality data and scanner information, or models for combinations of baseline-to-followup equipment settings, may improve the precision of longitudinal settings. The longitudinal model specified in (2) is linear, and could be expanded to nonlinear modeling cases in future studies. Finally, treatment and disease duration information were not available for this cohort, and future studies of more specialized clinical subpopulations promise further improvement for precision medicine applications.

The potential for growth charts for quantitative assessments of brain volumes and changes therein is vast. By leveraging longitudinal measurements in a statistically principled way, longitudinal growth charts that integrate image quality allow for precise and interpretable determinations of patients’ radiological changes in clinical practice settings. These may help to mitigate the clinicoradiological paradox of a dissociation between radiological findings and clinical outcomes (Barkhof, 2002) in management of the disease, and guide decisions about therapeutic changes. In research settings, GAMLSS will allow for more precisely charting of subject-specific changes and therapeutic effects. Future methodological studies on modeling of regionally-specific changes in brain volume and charting the evolution of lesion metrics are warranted, and the approach presented here can also be readily adapted beyond MS to other neurodegenerative diseases.

## Data/code availability statement

Due to privacy considerations with respect to patient consent and data sharing agreements, data are unfortunately not available for sharing. As the software developed is part of a proprietary product development effort, source code is not currently available for public distribution.

---

**Supplementary Figure 1:**
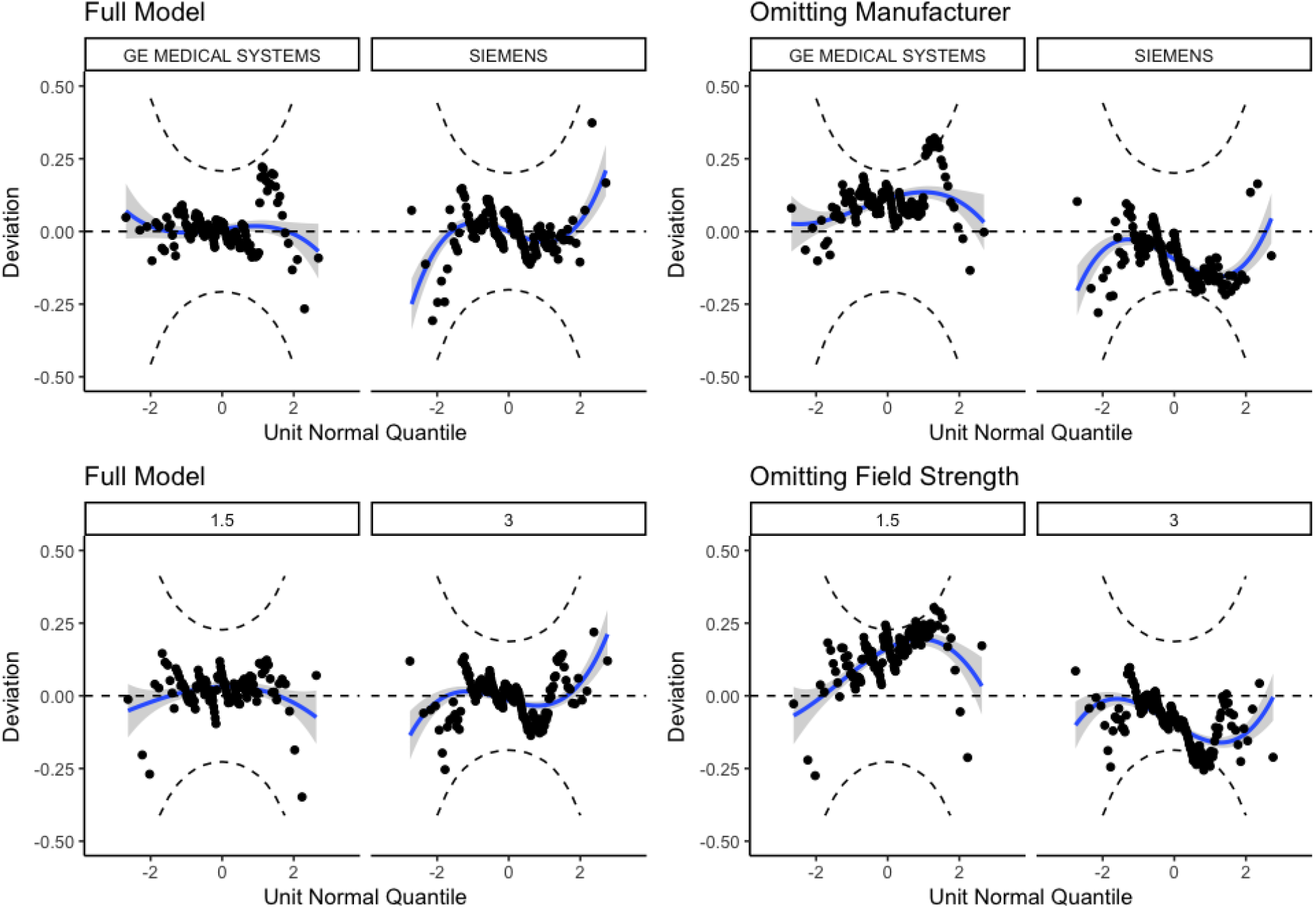
Illustrative example of worm plots from cross-sectional modeling of total brain volume in females. A worm plot is a detrended normal QQ-plot, which highlights deviations from normality in the model residuals. Worm plots where deviations are close to zero (horizontal dashed line) and within the confidence bands (inside the two elliptic dashed curves) indicate superior fit. Top row shows comparisons of the final model with scanner manufacturer in the model (left) versus without scanner manufacturer in the model (right) scanner manufacturer in the model, for data acquired on GE versus Siemens MRI systems. Bottom row shows comparisons of the final model with scanner field strength included in modeling (left) versus without scanner field strength included in modeling (right), for data acquired at 3T versus 1.5T field strength. Note that when scanner manufacturer is omitted (top right), there are deviations in average (e.g., the “worm” passes below the origin in Siemens data indicating that the fitted mean is too small), which are ameliorated by including manufacturer in the mean model (top left). When field strength is omitted (bottom right), there are deviations in average as well as slope (e.g., the worm has a positive slope in 1.5T data indicating that the fitted variance is too small), which are ameliorated by including mean and variance terms for field strength (bottom left).

